# Genomic and phenotypic comparisons reveal distinct variants of *Wolbachia* strain *w*AlbB

**DOI:** 10.1101/2022.02.25.482065

**Authors:** Julien Martinez, Perran A. Ross, Xinyue Gu, Thomas H. Ant, Shivan M. Murdochy, Lily Tong, Ana da Silva Filipe, Ary A. Hoffmann, Steven P. Sinkins

## Abstract

The intracellular bacterium *Wolbachia* inhibits virus replication and is being harnessed around the world to fight mosquito-borne diseases through releases of mosquitoes carrying the symbiont. *Wolbachia* strains vary in their ability to invade mosquito populations and suppress viruses in part due to differences in their density within the insect and associated fitness costs. Using whole-genome sequencing, we demonstrate the existence of two variants in *w*AlbB, a *Wolbachia* strain being released in natural populations of *Aedes aegypti* mosquitoes. The two variants display striking differences in genome architecture and gene content. Differences in the presence/absence of 49 genes between variants include genes located in prophage regions and others potentially involved in controlling the symbiont’s density. Importantly, we show that these genetic differences correlate with variation in *w*AlbB density and its tolerance to heat stress, suggesting that different *w*AlbB variants may be better suited for field deployment depending on local environmental conditions. Finally, we found that the *w*AlbB genome remained stable following its introduction in a Malaysian mosquito population. Our results highlight the need for further genomic and phenotypic characterization of *Wolbachia* strains in order to inform ongoing *Wolbachia*-based programmes and improve the selection of optimal strains in future field interventions.

**Importance:** Dengue is a viral disease transmitted by *Aedes* mosquitoes that threatens around half of the world population. Recent advances in dengue control involve the introduction of *Wolbachia* bacterial symbionts with antiviral properties into mosquito populations which can lead to dramatic decreases in the incidence of the disease. In light of these promising results, there is a crucial need to better understand the factors affecting the success of such strategies, in particular the choice of *Wolbachia* strain for field releases and the potential for evolutionary changes. Here we characterized two variants of a *Wolbachia* strain used for dengue control that differ at the genomic level and in their ability to replicate within the mosquito. We also found no evidence for the evolution of the symbiont within the two years following its deployment in Malaysia. Our results have implications for current and future *Wolbachia*-based health interventions.

## Introduction

*Aedes aegypti* mosquitoes are the primary vectors of dengue, a neglected viral disease ranked by WHO among the top ten global health threats, with 50–100 million clinically apparent cases and half a million hospitalizations for severe disease every year (1). Current control methods based on insecticide fogging for mosquito suppression have failed to halt the continued expansion in range and incidence of dengue, and rising levels of insecticide resistance mean that there is a pressing need for innovative approaches. *Wolbachia* are maternally inherited symbiotic bacteria found in many insect species, but not naturally in *Ae. aegypti* (Ross et al. 2020); however following lab transfer into this species some *Wolbachia* strains can efficiently block dengue transmission (3–6), by causing perturbations in various cellular pathways including lipid transport (7).

*Wolbachia* strains *w*Mel from *Drosophila melanogaster* and *w*AlbB from *Aedes albopictus* have both been shown to spread to and remain at a high frequency in *Ae. aegypti* populations following releases of *Ae. aegypti* at a comparatively modest scale and duration without needing continuous re-introduction (8–11). These strains have a self-spreading capability using a form of reproductive manipulation known as cytoplasmic incompatibility (CI), whereby the progeny of *Wolbachia*-carrying males and *Wolbachia*- free females die, while the reverse cross is fertile, giving an advantage to *Wolbachia*- carrying females. Both strains have been shown to efficiently reduce dengue transmission, providing a safe, sustainable, cost-effective and eco-friendly biocontrol tool that holds great promise for reducing the global burden of dengue (8, 11–14).

Since releases of *Ae. aegypti* carrying *w*AlbB (6) were carried out in Malaysia in sites around Kuala Lumpur that were previously hot-spots for dengue transmission, dengue has been substantially decreased (8). When larvae develop under high temperature regimes with diurnal peaks around 37°C, *w*AlbB is more stable than *w*Mel, maintaining a higher density, high maternal transmission and efficient dengue transmission blocking (6, 15–18). The fitness cost of *w*AlbB in *Ae. aegypti* is higher than *w*Mel in lab assays, mainly due to slightly reduced adult longevity (6) and reduced fertility and fecundity of adult females produced from quiescent eggs (19). *Wolbachia* fitness costs negatively affect population dynamics, raising the threshold frequency that must be exceeded for CI-mediated spread to occur (20–22) and for *Wolbachia* to remain at stable high frequency after introduction, as occurred with *w*AlbB at a number of sites in Malaysia (8). Several independent transinfections of *w*AlbB from *Ae. albopictus* have been generated in *Ae. aegypti* through microinjection (6, 23, 24) and two of these have been released in natural populations (8, 25). While the transinfections originate from different geographic locations, it is unclear if there are genetic or phenotypic differences between them.

The effectiveness of *Wolbachia* interventions against dengue could be compromised in the longer term by evolutionary changes in the *Wolbachia* or mosquito genome (26). Virus transmission blocking could be reduced over time if mosquito-*Wolbachia* co-evolution results in lower *Wolbachia* density overall, or more restricted tissue distribution to the ovaries and testes. The *w*AlbB-associated reduced hatch rate of stored *Ae. aegypti* eggs could also be ameliorated by natural selection (27); if this selection acts specifically at the egg stage and does not impact the dengue transmission-blocking phenotype, it would be advantageous overall for implementation of the strategy. No obvious phenotypic changes have been observed in *w*AlbB to date in field populations of *Ae. aegypti* (18) but longer-term monitoring is required.

The primary aim of this study was to sequence the genome of the *w*AlbB strain released in Malaysia. This is useful for several reasons: to be able to ascertain whether this *w*AlbB has any unique genomic features relative to previously published *w*AlbB genomes; to be able to track genomic evolution that may occur in the field, that could potentially compromise the effectiveness of the dengue intervention; and to allow for the creation of molecular assays to allow this variant to be distinguished from the naturally occurring *w*AlbB present in *Ae. albopictus*, which will be useful in *Wolbachia* frequency monitoring, since both species are present in the intervention sites. Other aims were to compare the impact of different *w*AlbB infections on *Wolbachia* density, egg quiescence and responses to heat.

## Results

### Comparative genomics of *w*AlbB genomes

We compared three publicly available *w*AlbB circular genomes (Texas, Florida and Hainan) with a new draft assembly that we generated from an Indonesian *w*AlbB infection previously transferred into *Ae. aegypti* (6) (Figure 1A). The Indonesian *w*AlbB assembly was 1.45 Mb in size, >99% of which was made of 9 contigs (Table 1). This is slightly shorter than finished *w*AlbB genomes (1.48 Mb), however, similar numbers of single-copy conserved orthologues were found between *w*AlbB genomes suggesting the *w*AlbB-II assembly is nearly complete (Table 1). In light of the genomic differences described below, we will refer to *w*AlbB-I and *w*AlbB-II to designate the reference variant from Texas and the Indonesian variant respectively.

**Figure 1.**
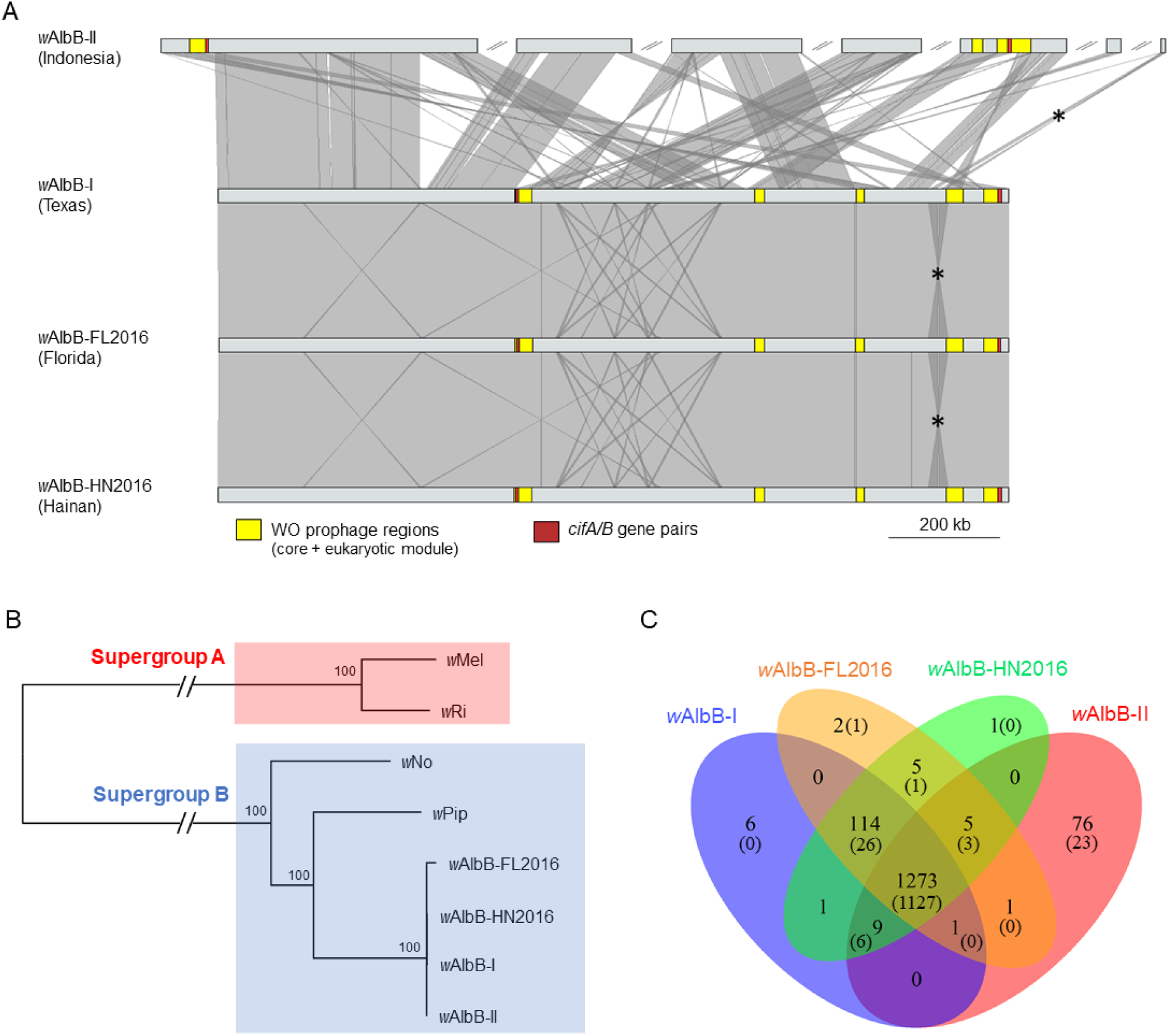
Comparative analysis and phylogeny of *w*AlbB genomes. (A) genome-wide synteny. Grey areas between genomes indicate similarities based on a megablastn comparison. * indicates a seven-gene region that is duplicated in *w*AlbB-I, -FL2016 and - HN2016. Blast hits and contigs < 3,000 bp were excluded from the figure and some contigs were reoriented to improve visualization. (B) Maximum likelihood phylogeny using a concatenated alignment of 614 orthologous genes. Node labels are bootstrap supports calculated from 1,000 replications. (C) Venn diagram showing numbers of orthologs shared between *w*AlbB genomes (numbers in parentheses exclude transposable elements).

**Table 1.** Genome features of sequenced wAlbB genomes.

|  | wAlbB-II | wAlbB-I | wAlbB-FL2016 | wAlbB-HN2016 |
| --- | --- | --- | --- | --- |
| Genbank accession | JAKLOR00000000 | CP031221.1 | CP041923.1 | CP041924.1 |
| Geographical origin | Indonesia | Texas | Florida | Hainan |
| Assembly size (bp) | 1,450,135 | 1,484,007 | 1,482,279 | 1,483,853 |
| Contigs (>1000 bp) | 16 (9) | 1 | 1 | 1 |
| GC content (%) | 34.3 | 34.4 | 34.4 | 34.4 |
| gene | 1,365 | 1,434 | 1,426 | 1,428 |
| CDS | 1,324 | 1,393 | 1,385 | 1,387 |
| rRNA | 3 | 3 | 3 | 3 |
| tRNA | 34 | 34 | 34 | 34 |
| BUSCO complete<sup>a</sup> | 328 | 328 | 326 | 328 |
| BUSCO fragment<sup>a</sup> | 9 | 9 | 11 | 9 |
| BUSCO missing<sup>a</sup> | 95 | 95 | 95 | 95 |

<sup>a</sup>The BUSCO analysis was run against the alphaproteobacteria\_odb10 database.

The Indonesian *w*AlbB-II variant clusters with the three other available *w*AlbB genomes into a monophyletic clade within *Wolbachia* supergroup B (Figure 1B). However, despite the strong phylogenetic relatedness, *w*AlbB-II displays striking differences in genome synteny when compared to *w*AlbB-I and the *w*AlbB-I-like genomes originating from Florida and China (Figure 1A). There are three incomplete WO prophage regions in the *w*AlbB-II genome, indicating ancient WO phage infections. The structural and non-structural modules (head, tail, baseplate, replication and eukaryotic modules) are split between the different regions, suggesting no active phage replication (Figure 2). Moreover, essential genes of the phage head, tail and baseplate are missing indicating that the production of phage particles is impaired (Table 2). Core phage genes are present in single copies except for the recombinase and phospholipase D which are both present in two copies with high sequence divergence, suggesting that the *w*AlbB genome may have been colonized in the past by more than one WO phage (Table 2). The other *w*AlbB genomes also carry prophage regions with sequence similarities to those of *w*AlbB-II but these have rearranged into five different regions (Figure 1A). Two pairs of the cytoplasmic incompatibility-inducing genes *cifA* and *cifB* are located within *w*AlbB-II’s prophage regions and are identical to the other *w*AlbB genomes. The two gene pairs are related to type III *cif* homologues for one pair and type IV for the other pair as defined in previous studies (28, 29).

**Figure 2.**
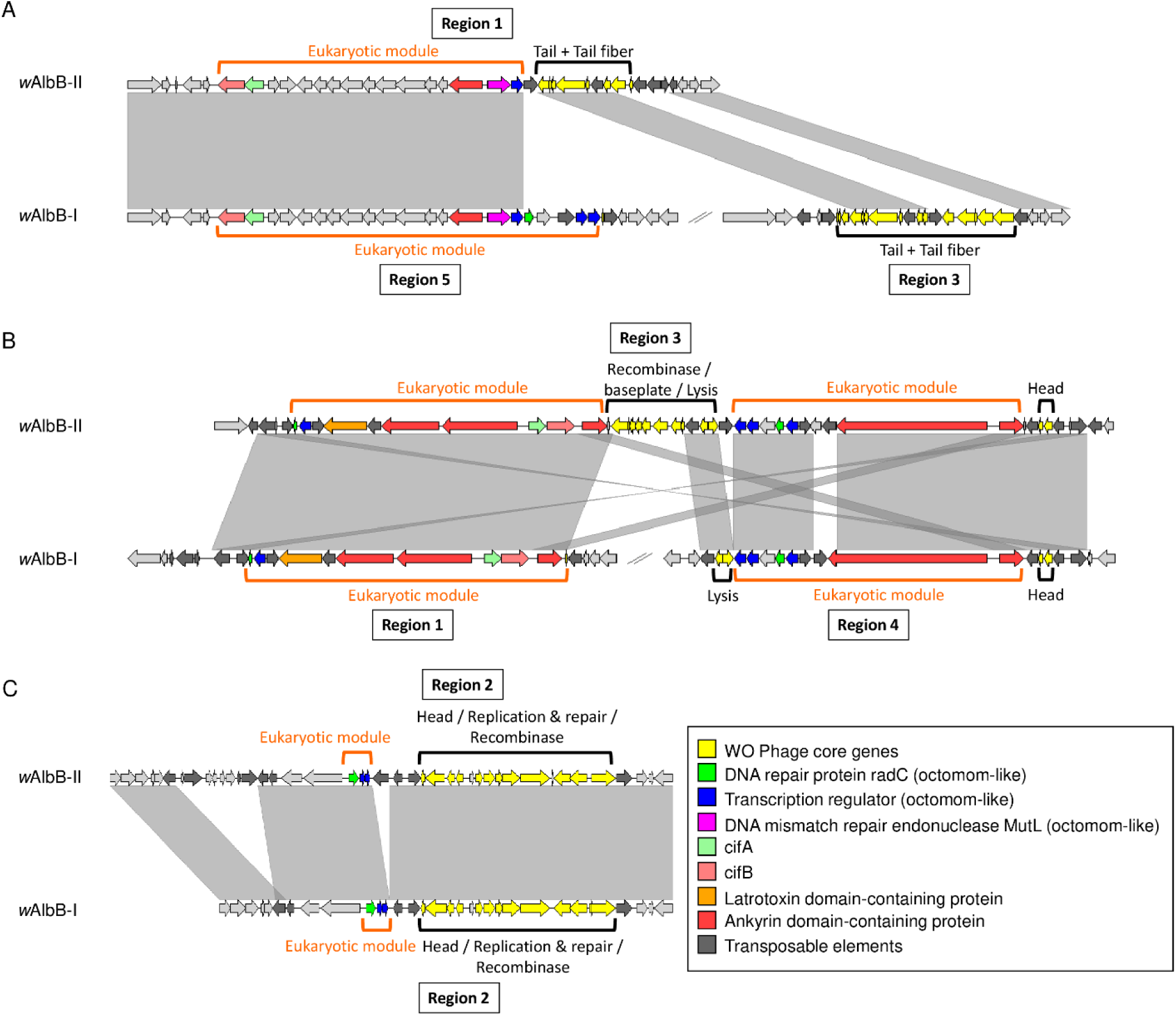
Synteny of WO prophage regions between *w*AlbB-I and *w*AlbB-II genomes. Grey areas indicate similarities based on megablastn comparisons. Blast hits < 2,000 bp were excluded from the figure to improve visualization. Panels depict prophage region 1 (A), region 2 (B) and region 3 (C) in the *w*AlbB-II assembly.

**Table 2.**
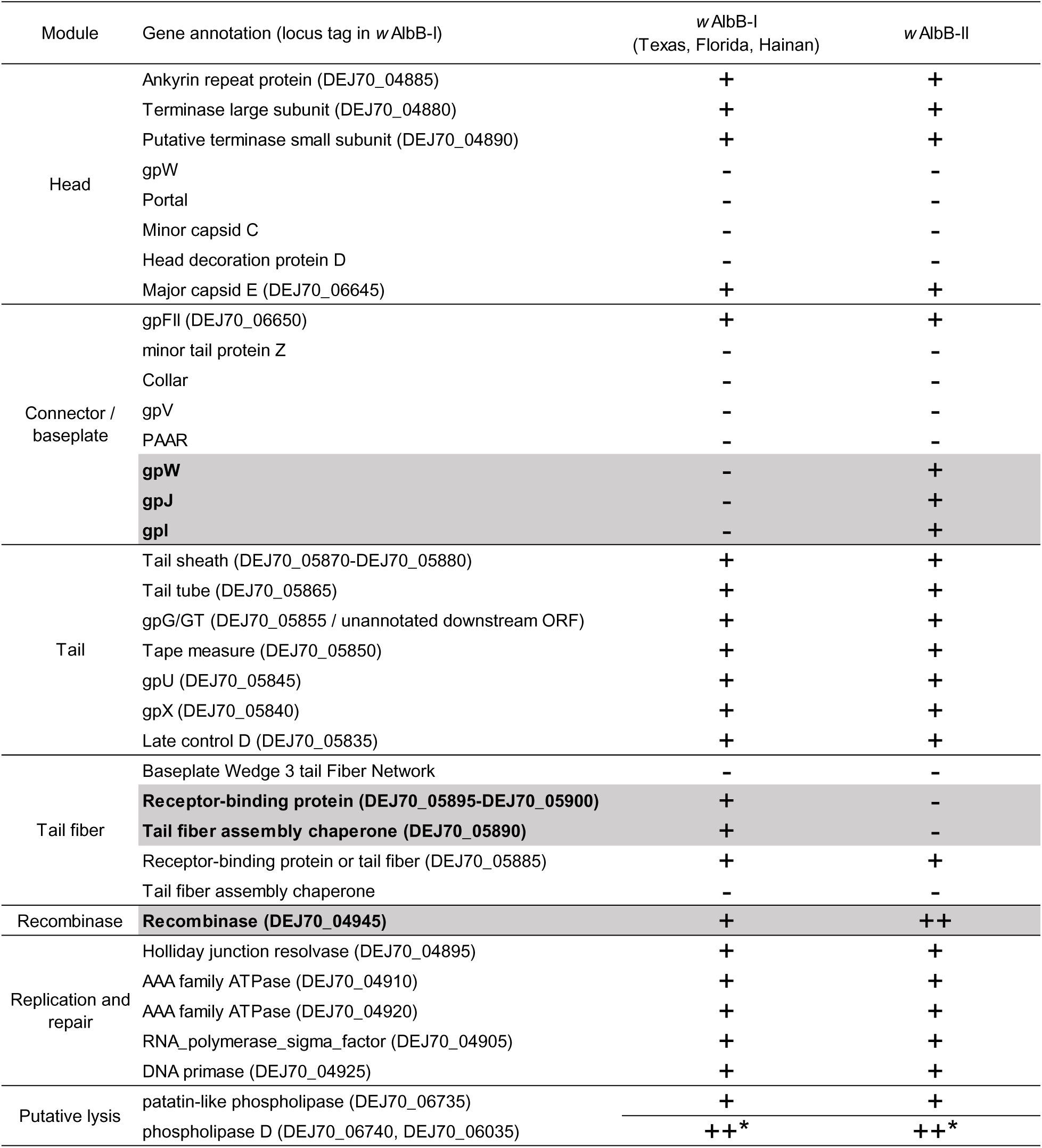
Presence/absence of WO phage core genes in prophage regions. Gene is either present (+) or absent (-). Differences between variants are highlighted in grey. *one copy of the phospholipase D is located outside prophage regions.

In addition to chromosomal rearrangements, we found noticeable differences in gene content between the *w*AlbB-I-like and *w*AlbB-II genomes (Figure 1C, Table S1), with ∼70% of the differences involving repeat elements (transposases, group II introns reverse-transcriptase and related pseudogenes). Excluding repeat elements, *w*AlbB-II harbours 23 genes that are absent from or pseudogenized in the three other genomes, while on the other hand, *w*AlbB-I, -FL2016 and -HN2016 share 26 genes not found or pseudogenized in the *w*AlbB-II draft assembly. Some of this variation is located within and around prophage regions, where *w*AlbB-II and the other *w*AlbB genomes have lost different core and accessory phage genes (Figure 2, Table 2, Table S1). For instance, one of the *w*AlbB- II phage eukaryotic modules lacks two copies of a putative transcriptional regulator and one copy of a DNA repair protein which are homologues of genes in an eight-gene locus know as Octomom thought to influence *Wolbachia* proliferation in *w*Mel-like strains (30–32) (Figure 2A). Interestingly, *w*AlbB-II also carries two syntenic proteins (WP_019236968.1 and WP_019236969.1) with homologies to arthropod protein translocase subunit secA genes that are absent in other *w*AlbB genomes (Table S1) but present in other *Wolbachia* strains. The two genes are of unusual length for *Wolbachia* genes. They branch with a few other *Wolbachia* homologues within arthropod lineages with no closely-related bacterial homologues indicating two independent horizontal transfers from arthropods to *Wolbachia* in the case of WP_019236968.1 and at least one such event for WP_019236969.1 (Figures S2). Finally, several genes differed between *w*AlbB variants due to pseudogenization by the insertion of a transposon. For example, a homologue of the *Wolbachia* surface protein *wspB*, is pseudogenized in the three *w*AlbB- I-like genomes while a full-length version of the gene is present in *w*AlbB-II.

It is possible that some of the genes predicted to be absent in *w*AlbB-II might in fact be present but were not assembled. For instance, one of *w*AlbB-II contigs showed a 2x sequencing depth compared with the rest of the assembly and is identical to a large region that is duplicated in the other *w*AlbB genomes (asterisks in Figure 1A, Table S1). Similarly, another three-gene duplication in *w*AlbB-I genomes also displayed a 2x sequencing depth in *w*AlbB-II (Table S1). Thus, we counted the extra copies of genes in these two large duplications as present in *w*AlbB-II. Nevertheless, we confirmed that all other genes missing in *w*AlbB-II (excluding transposable elements) were either present but annotated as pseudogenes or that no *w*AlbB-II Illumina reads mapped to the corresponding genes in the other genomes (Table S1, Figure S1).

Numerous single nucleotide polymorphisms (SNPs) and small indels between the *w*AlbB genomes were identified (Table 3, Table S2). In reciprocal comparisons, there were more polymorphisms when *w*AlbB-II was used as the reference genome for SNP calling. This is likely because the *w*AlbB-II assembly is incomplete, which may generate false-positives within misassembled repeat sequences. This is supported by the fact that five SNPs were detected in transposable elements when mapping *w*AlbB-II reads on its own assembly.

**Table 3.** Summary of the SNP analysis.

|  | Genome used as reference for SNP calling |  |  |  |
| --- | --- | --- | --- | --- |
| Illumina reads | wAlbB-II | wAlbB-I | wAlbB-FL2016 | wAlbB-HN2016 |
| wAlbB-II | 5 | 177 | 793<sup>a</sup> | 318<sup>a</sup> |
| wAlbB-MC | 13 | 195 | 801<sup>a</sup> | 343<sup>a</sup> |
| wAlbB-I | 201 | 0 | 930<sup>a</sup> | 361<sup>a</sup> |

<sup>a</sup>A high proportion of the SNPs against wAlbB-FL2016 and -HN2016 show an unusual distribution and may be sequencing errors in the reference genome (Table S2).

Moreover, the five SNPs were all polymorphic upon visual inspection of the mapped reads. Nevertheless, *w*AlbB-II showed the smallest number of SNPs against *w*AlbB-I, while it was more divergent from *w*AlbB-FL2016 and *w*AlbB-HN2016. This is inconsistent with the pattern of gene presence/absence observed above. However, a large proportion of SNPs against *w*AlbB-FL2016 and *w*AlbB-HN2016 displayed an atypical distribution with a few loci accumulating the majority of the SNPs. As noted in Ross et al. (2021), this could be due to possible DNA contamination in these two assemblies. Around 60% of the SNPs between *w*AlbB-II and *w*AlbB-I are non-synonymous, of which some are located within ankyrin repeat domain-containing genes as well as genes potentially involved in transcription and RNA processing (e.g. sigma factor RpoD (34), transcription elongation factor NusA (35), RNA polymerase subunit alpha and beta, ribonucleases E and D (36)), protein synthesis (e.g. ribosomal proteins, translational GTPase TypA (37)), cell wall synthesis and remodelling (e.g. N-acetylmuramoyl-L-alanine amidase, D-alanyl-D-alanine carboxypeptidase, M23 family peptidase, UDP-N-acetylmuramate dehydrogenase (38, 39)) and stress response (e.g. heat shock proteins: ATP-dependent Clp endopeptidase (40), DegQ endoprotease (41)).

### *w*AlbB *Wolbachia* variants display background-dependent differences in density

To determine potential effects of *w*AlbB variation on phenotype, we generated *Ae. aegypti* populations with different *Wolbachia* infection types (*w*AlbB-I, *w*AlbB-II or uninfected) and backgrounds (Australian (Au) or Malaysian (My)) through reciprocal backcrossing (33). *Wolbachia* density was influenced by *w*AlbB variant in both sexes (Table S3), with *w*AlbB-I individuals having higher *Wolbachia* densities than *w*AlbB-II individuals (Figure 3A and B). We found no clear effect of nuclear background for either sex, but there was a significant interaction between nuclear background and *w*AlbB variant in males (Table S3). In the Australian background, *w*AlbB-I had a higher density than *w*AlbB-II (GLM: females: F_1,76_ = 13.792, P < 0.001, males: F_1,73_ = 27.401, P < 0.001) but there were no significant differences between variants in the Malaysian background (females: F_1,76_ = 0.982, P = 0.325, males: F_1,76_ = 2.225, P = 0.140).

**Figure 3.**
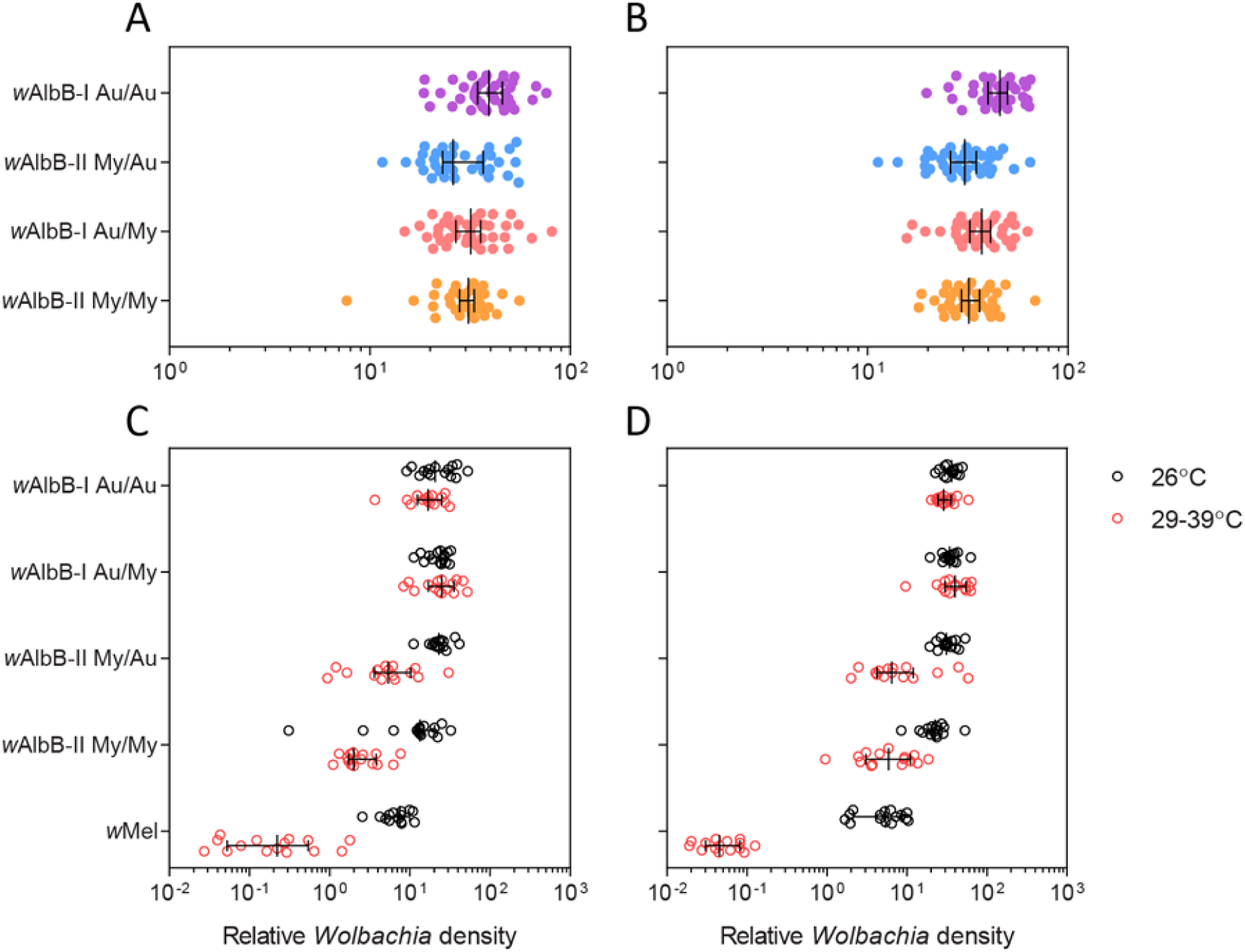
Differences in density between *w*AlbB variants. Female (A) and male (B) *Wolbachia* density in reciprocally-backcrossed *Aedes aegypti* populations. Populations have different combinations of *Wolbachia* infection type/mitochondrial haplotype (*w*AlbB- I and Au or *w*AlbB-II and My) and nuclear background (Au or My). Data from two replicate populations were pooled for visualization. *Wolbachia* density in (C) females and (D) males following exposure to cyclical heat stress during the egg stage. Eggs were exposed to cyclical temperatures of 29-39°C for 7 d (red circles) or held at 26°C (black circles). Each point represents the relative density for an individual averaged across 2-3 technical replicates. Medians and 95% confidence intervals are shown in black lines. Data for *w*AlbB-I and *w*Mel have also been included from (33).

To test the stability of *w*AlbB variants at high temperatures, we measured *Wolbachia* densities in adults after eggs were exposed to cyclical heat stress (29-39°C) or held at 26°C for one week. *Wolbachia* density was influenced by *w*AlbB variant and temperature treatment, with significant interactions between *w*AlbB variant and nuclear background as well as *Wolbachia* variant and temperature (Table S4). When the *w*AlbB-I and *w*AlbB-II variants were tested separately, we found no effect of temperature or nuclear background in either sex (all P > 0.141) for *w*AlbB-I, indicating that this infection is stable under heat stress (Figure 3C and D). In contrast, *w*AlbB-II density was lower in the heat stress treatment (Females: F_1,56_ = 51.940, P < 0.001, Males: F_1,56_ = 61.814, P < 0.001) and in the Malaysian background (Females: F_1,56_ = 12.831, P = 0.001, Males: F_1,56_ = 4.549, P = 0.037). Across both sexes and backgrounds, median *w*AlbB-II density under cyclical heat stress decreased by 80.4%, compared to 8.1% for *w*AlbB-I and 98.7% for *w*Mel.

### Quiescent egg viability depends on mosquito nuclear background and *w*AlbB infection

Stored eggs from populations with different combinations of *w*AlbB infection type (*w*AlbB- I, *w*AlbB-II or uninfected), mitochondrial haplotype (Au or My) and background (Au or My) were hatched every three weeks to determine quiescent egg viability. *w*AlbB infection greatly reduced quiescent egg viability in all four combinations of background and mitochondrial haplotype (Figure 4). By week 16, hatch proportions for *w*AlbB-infected populations approached zero while hatch proportions for uninfected populations exceeded 40%. In uninfected populations, we found significant effects of egg storage duration and nuclear background on egg hatch proportions (Table S5). Eggs with an Australian background had higher hatch proportions (median 0.823) than eggs with a Malaysian background (median 0.503) by the end of the experiment. In *w*AlbB-infected populations, we found significant effects of egg storage duration, squared egg storage duration, nuclear background and replicate population (Table S5). Although populations carrying *w*AlbB-II had higher overall hatch proportions than *w*AlbB-I populations in both backgrounds (Figure 4), effects of *w*AlbB variant were not significant (Table S5). We had low power to detect *w*AlbB variant effects in this analysis due to nesting replicate population within *w*AlbB variant.

**Figure 4.**
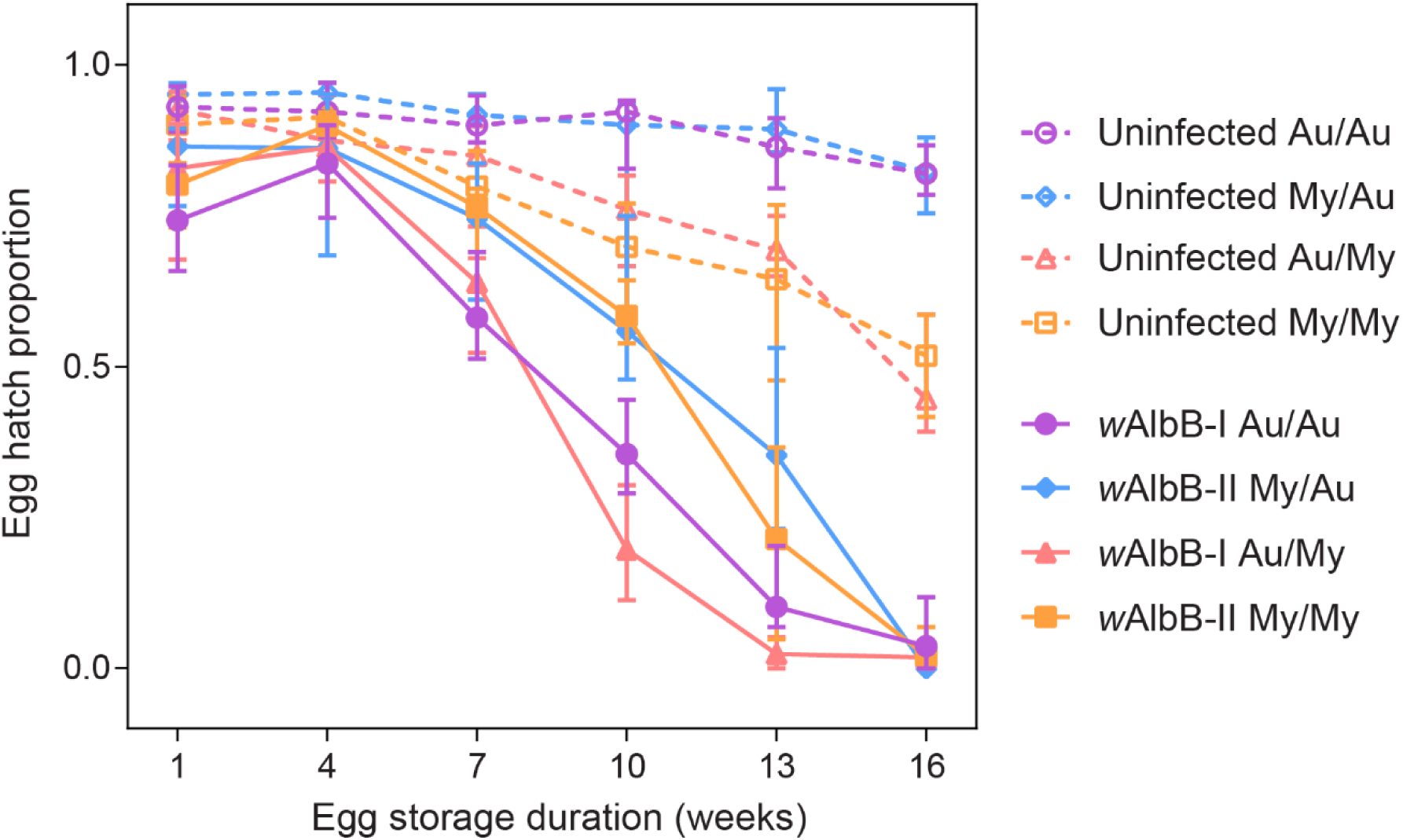
Quiescent egg viability of reciprocally-backcrossed *Aedes aegypti* populations. Populations have different combinations of *Wolbachia* infection type (*w*AlbB-I, *w*AlbB-II or uninfected), mitochondrial haplotype (Au or My) and nuclear background (Au or My). Data from two replicate populations were pooled for visualization. Symbols show median egg hatch proportions while error bars show 95% confidence intervals. Data for Au/Au and Au/My populations have also been included from (33).

### A multiplex PCR reaction for diagnostics of *w*AlbB variants

Given the genomic differences observed between the *w*AlbB genomes, we developed a PCR reaction allowing the distinction between the *w*AlbB-I and -II variants in *Ae. Aegypti* and *Ae. albopictus*. From our gene content analysis, we selected an AAA-family ATPase protein and the phage tail formation protein I as markers to distinguish the two *w*AlbB variants. Primers for these markers were designed and pooled with primers amplifying the 18S mosquito control gene into a multiplex PCR reaction. *w*AlbB-I in the Aa23 *albopictus* cell line and *w*AlbB-II in our *Ae. aegypti* mosquito lab line displayed the expected band profiles and were clearly distinguishable on the agarose gel (Figure 5). Moreover, the *w*AlbB infecting a Malaysian line of *Ae. albopictus* mosquitoes (JF) displayed identical bands to *w*AlbB-I variant.

**Figure 5.**
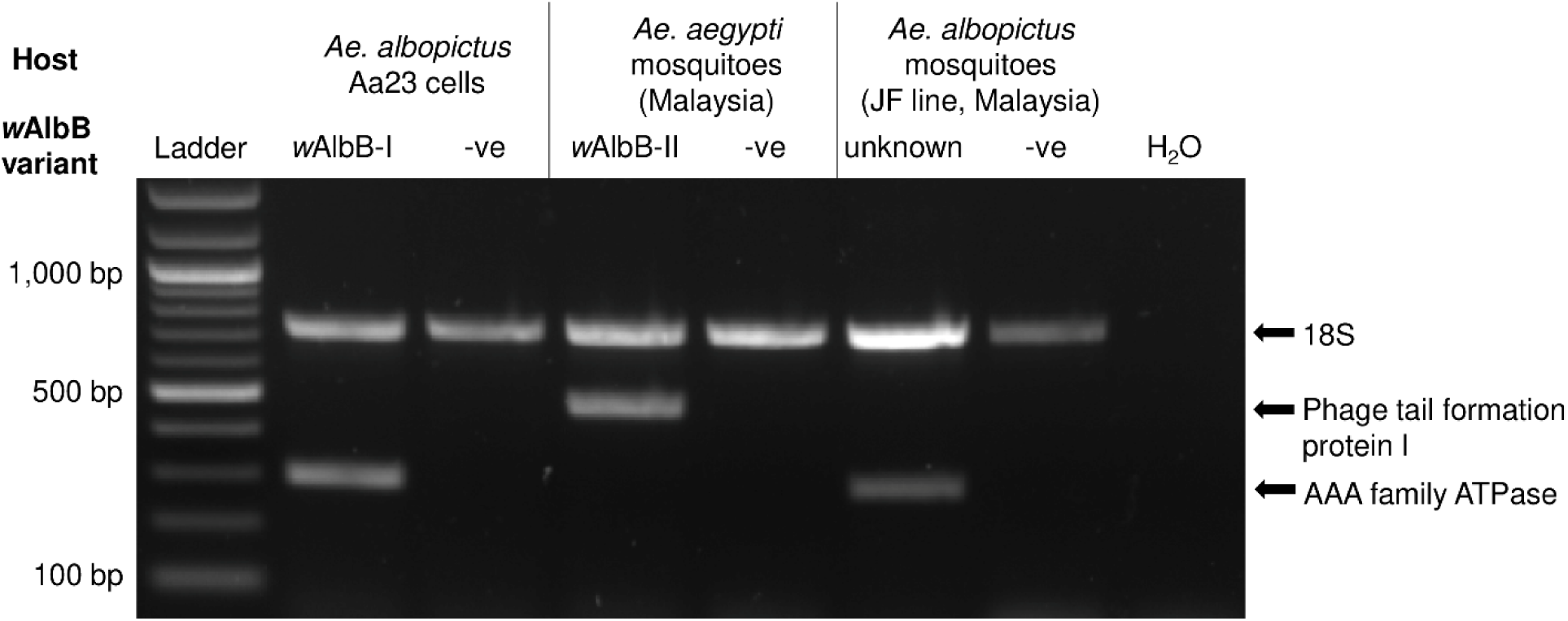
*w*AlbB variant-specific multiplex PCR reaction. DNA was extracted from approximately 1.10^6^ cells and five mosquito females for the Aa23 and mosquito samples respectively. -ve: *Wolbachia*-negative controls.

### No evidence for evolution of *w*AlbB-II genome following field release

Using the *w*AlbB-II assembly as a reference, we mapped the sequencing data generated from a *w*AlbB-infected *Ae. aegypti* colony *w*AlbB-MC (Mentari Court site) that was isolated from the field two years following field releases of wAlbB-II in Malaysia (18). All genome positions of the reference assembly were covered by read data from *w*AlbB-MC with no drastic drop in sequencing depth suggesting that no gene was lost since field releases (Figure S1A). Thirteen SNPs were detected, however, all were located within transposable elements (Table 3 and Table S2). Similar to *w*AlbB-II sequencing data, visual inspection of the *w*AlbB-MC mapped reads revealed that all SNPs were highly polymorphic indicating that they are likely false-positives caused by the use of an incomplete reference genome. Therefore, we conclude that there is little evidence for genomic changes that may have occurred following the introduction of *w*AlbB-II in the field.

## Discussion

Here we uncovered major genomic differences between closely-related *w*AlbB-strain *Wolbachia* and showed these differences correlate with variation in symbiont density. *w*AlbB diversity has commonly been investigated using a limited number of markers such as the *wsp* gene and Multi-Locus Sequence Typing (MLST) genes with little to no variation observed between isolates from different locations (42–44). Whole-genome sequencing provides a higher resolution and has revealed significant genomic differences between closely-related *Wolbachia* strains (45–47). Using this method, we demonstrated the existence of at least two types of *w*AlbB variants that differ in genome synteny and gene content. The reference *w*AlbB-I variant originating from Texas and the *w*AlbB isolates from Florida and China share a similar genome architecture and gene content while the Indonesian *w*AlbB-II variant is strikingly different. Further sampling of *w*AlbB genomic diversity will provide new insights into the evolution of this symbiont lineage and may help unravel the colonization history of its native host, *Ae. albopictus*, across continents (48).

Importantly, there are differences in symbiont density between the *w*AlbB-I and *w*AlbB-II variants. Although only one isolate representative of each variant was characterised here, these results suggest that variant-specific genetic determinants may be driving some of the differences in density. It is possible, but less likely, that differences in mitochondrial haplotypes contributed to some of the variation in symbiont density since mitotypes were not cross-factored with the *w*AlbB variants. The reference isolates for *w*AlbB-I and II variants differed in the presence/absence of 49 genes (excluding transposable elements) and on ∼200 SNPs, 60% of which were non-synonymous. How much of this variation contributes to differences in symbiont density remains to be investigated, and further phenotypic characterisation of new variants will help shorten the list of candidate genes since there is currently no transformation system in *Wolbachia* for functional validation. Interestingly, some of the genomic differences are located within prophage regions. It has been hypothesised that WO phage replication lowers *Wolbachia* density (49); however, we only found incomplete prophage regions, suggesting that active phage mobilisation is unlikely to occur in *w*AlbB. Alternatively, variation in the expression of phage accessory genes could be responsible for differences in density. Indeed, *w*AlbB-I carries additional copies of a DNA repair protein *radC* and a putative transcriptional regulator in its phage eukaryotic module. Homologues of these genes in *w*Mel-like strains, WD0507 and WD0508, are part of a 20 kb region called Octomom. Octomom is to date the only genetic determinant shown to influence *Wolbachia* proliferation in a copy-number dependent manner (30). Both complete loss of the region and amplification have been associated with symbiont over-proliferation, which in turn has negative effects on host lifespan (31, 32).

Density variation could also stem from the way *w*AlbB variants interact with the host. *Wolbachia* genomes commonly harbour an array of ankyrin domain-containing genes which are predicted to be involved in protein-protein interactions as well as secretion systems that may allow the export of bacterial effectors into the host cell cytoplasm (50, 51). Several ankyrin domain-containing genes are among the candidate genes that differ between *w*AlbB variants. Additionally, *w*AlbB-II harbours two syntenic proteins showing homologies to arthropod protein translocase subunit SecA genes. Homologues were previously found in other *Wolbachia* strains (52) and their phylogenetic distribution points towards an acquisition through horizontal transfer from a eukaryotic host. SecA proteins are involved in the transport of bacterial and ER-exported proteins (53), suggesting that the *w*AlbB variants may differ in the way they interact with the host cell. Interestingly, *w*AlbB variants also differed in a homologue of the *Wolbachia* surface protein *wspB*. *wspB* is pseudogenized in *w*AlbB-I and this variant maintains higher densities under a high temperature cycle compared to *w*AlbB-II that carry a full-length version of the gene. This is consistent with two recent studies showing that, among variants of the *Wolbachia* strain *w*Mel, pseudogenization of the *wspB* gene is associated with variation in symbiont density and maternal transmission and that the magnitude of this effect can vary with both temperature and host background (54, 55). Finally, *w*AlbB variants differed in a number of house-keeping genes involved in essential functions such as DNA replication, RNA processing, translation or cell wall biogenesis that may contribute to the variation in symbiont density, and in heat shock response genes that could potentially control their different degrees of tolerance of heat stress.

The phenotypic differences detected impact on the relative ability of these variants to spread through *Ae. aegypti* populations, and their efficacy for dengue control programmes. The declining hatch rates over time of dried quiescent eggs is an important component of the fitness costs of *w*AlbB in this host (19). In areas where a high proportion of larval sites are temporary and experience intermittent inundation, dry eggs are often in quiescence for extended periods, and thus the fitness cost of *w*AlbB will be higher relative to *Wolbachia*-free wildtype counterparts than when larvae develop in permanent breeding sites such as water storage tanks. This factor will increase the threshold population frequency that must be exceeded for *Wolbachia* to spread / remain stable in the population. The higher, background-dependent negative impact of the *w*AlbB-I on quiescent egg hatch rates means that this variant will be predicted under some backgrounds / ecological conditions to spread less efficiently than *w*AlbB-II. Conversely the apparent slightly higher tolerance of *w*AlbB-I for very high temperature egg storage may also impact on relative spread dynamics, since maintenance of higher density of *w*AlbB-I may ultimately impact maternal transmission rates and virus inhibition. In the hottest climates in which *Ae. aegypti* occurs, *w*AlbB-I may prove to be a better option for dengue control than *w*AlbB-II. However, more data is needed under a variety of conditions and over multiple generations and life stages to test the relative impacts of temperature in more detail.

Previously, we found negligible changes in the *w*AlbB-I genome following transfer from its native host *Ae. albopictus* to *Ae. aegypti* (33) suggesting that there is little selective pressure to adapt to this new host, at least in laboratory conditions. This is in line with little genomic changes observed following artificial transfers of multiple *Wolbachia* strains between *Drosophila* species (56). Here we found no evidence of wAlbB-II genome evolution after its introduction in a field population of *Ae. aegypti*, which supports our earlier results showing stable density and antiviral effects using the same field-caught mosquito colony (18). It is also in line with the observed *w*Mel genome stability following the introduction of *Wolbachia*-infected *Ae. aegypti* mosquitoes in Australia (57, 58). *w*Mel and *w*MelPop-CLA infections in *Ae. aegypti* also show few long-term phenotypic changes following transinfection (10, 59).

## Methods

### *Wolbachia* purification

In order to generate Illumina sequencing data for both the reference *w*AlbB-II genome and its field-caught counterpart *w*AlbB-MC, *Wolbachia* was purified from whole mosquitoes. *w*AlbB-MC-infected mosquitoes were used after three generations spent in the lab since field collection in Mentari Court. For each genome, around 400 mosquitoes were collected into a 50 ml Falcon tube and snapped at −20°C for 10 min. Mosquitoes were then surface-sterilized for 3 min in 50% bleach, followed by 3 min in 70% ethanol and rinsed 3 times with sterile water. Mosquitoes were then manually homogenized with 3 mm glass beads in 40 ml of Schneider’s media by shaking and further homogenized with a tissue-lyzer after transferring the homogenate into 2 ml tubes with 1 mm beads. Homogenates were centrifuged at 2,000 g for 2 min to remove tissue debris and the supernatant was sequentially filtered through 5, 2.7 and 1.5 µm sterile filters. The filtrate was aliquoted in Eppendorf tubes and centrifuged at 18,500 g for 15 min to pellet bacteria. The supernatant was discarded, Schneider’s media added and the previous centrifugation step repeated once. The bacterial pellet was resuspended in Schneider’s media and treated with DNase I at 37°C for 30 min to remove host DNA. Following digestion, samples were centrifuged at 18,500 g, the supernatant discarded and the DNase inactivated at 75°C for 10 min. Finally, bacterial pellets were pooled into one tube and DNA extracted with the Gentra Puregene tissue kit (Qiagen) with resuspension of the DNA pellet in 100 µl of nuclease-free water.

For long-read sequencing of the *w*AlbB-II genome, the same protocol was followed except that *Wolbachia* was purified from 150 freshly laid mosquito eggs. In order to increase the amount of starting DNA needed for Nanopore library construction, the DNA was amplified directly from the bacterial pellet before the DNA extraction step using the REPLI-G Midi kit (Qiagen).

### Whole-genome sequencing and genome assembly

For both *w*AlbB-II and *w*AlbB-MC, DNA libraries were prepared using the Kapa LTP Library Preparation Kit (KAPA Biosystems, Roche7961880001) and sequenced on the Illumina MiSeq platform with the MiSeq Reagent Kit v3 to generate 2×150 bp reads. Raw reads were demultiplexed using bcl2fastq and adapters were trimmed with Trimmomatic v0.38.0 (60). To generate long reads for *w*AlbB-II, 1 µg of whole-genome amplified gDNA was sheared into ∼8 kb fragments followed by purification and size-selection using AMPure XP beads (Beckman Coulter). Oxford Nanopore Technology (ONT) sequencing library preparation was then carried out with the Ligation Sequencing Kit (SQK-LSK109) and the library loaded on a MinION Flow Cell and sequenced for 72 hours using a GridION (ONT) controlled by MinKNOW software v20.06.9. Base-calling and demultiplexing were performed within MinKNOW using Guppy v4.0.11. ONT adapters were removed with Porechop v0.2.4 (61). Host reads were filtered out by mapping the Illumina and Nanopore reads against the *Ae. aegypti* reference assembly (Genbank accession: GCF_002204515.2) using Bowtie2 v2.4.2 (62) and Minimap2 v2.23 (63) respectively. Unmapped reads were then assembled using the Unicycler hybrid assembly pipeline (64). Contigs were visualized and blasted against several *Wolbachia* genomes in Bandage (65) and non-*Wolbachia* sequences were discarded from the assembly.

### Comparative genomics

The *w*AlbB-II assembly was compared to three other *w*AlbB isolates for which genome sequences are publicly available – the reference isolate (*w*AlbB-I), which is derived from *Ae. albopictus* mosquitoes caught in Houston, Texas, USA in 1986 and has subsequently been maintained in the *Ae. albopictus* Aa23 cell line (66, 67); an isolate from *Ae. albopictus* caught in St. Augustine, Florida, USA in 2016 (*w*AlbB-FL2016); and an isolate from *Ae. albopictus* caught in Haikou, Hainan, China in 2016 (*w*AlbB-HN2016). *w*AlbB genomes were all reannotated using Prokka v1.14.6 (68) prior to the gene content analysis. Roary v3.13.0 (69) was then used to determine the core and accessory genomes with a 95% identity threshold. The Roary output was manually curated to fix issues with pseudogenes in the accessory genomes by visualizing the genome annotations in the Artemis genome browser v16.0.0 (70). Large indels were confirmed by visual inspection of sequencing depth after mapping the *w*AlbB-II and *w*AlbB-I (SRA accession: SRR7623731) Illumina reads onto the different *w*AlbB genomes with Bowtie2 (Figure S1). Tblastn searches were run to locate WO phage genes, including the homologues of the Octomom and *cif* genes within the genomes using representative sequences. Prophage regions and their eukaryotic modules were then manually re-annotated by following the most recent guidelines (71). Whole-genome and prophage regions synteny were visualized using the R package genoPlotR (72). The maximum likelihood phylogenetic tree was inferred with RaxML v7.7.6 (73) using a core gene alignment of several Supergroup A and B *Wolbachia* genomes generated by Roary.

The SNP analysis was conducted using the Snippy pipeline v4.6.0 (Seeman, Torsten. Snippy: fast bacterial variant calling from NGS reads; 2020 https://github.com/tseemann/snippy). Illumina reads were mapped onto the different *w*AlbB reference genomes and SNPs were called with minimum mapping quality of 20, 10 reads minimum coverage and 0.9 minimum proportion for variant evidence. From the alignment BAM files, sequencing depth was calculated using Samtools depth v1.13 and visually inspected in R and Artemis to identify large indels and duplicated regions.

### Origin of *w*AlbB variants and mosquitoes used in phenotypic assays

The *w*AlbB-I variant used in phenotypic comparisons originates from *Ae. aegypti* mosquitoes that were transinfected in 2005 (24) with the same *w*AlbB infection as the reference isolate found in the Aa23 *Ae. albopictus* cell line. *w*AlbB-I was then transinfected into an *Ae. aegypti* line with an Australian mitochondrial haplotype (33). The *w*AlbB-I genome is almost identical to the *w*AlbB reference genome differing by only four Single Nucleotide Variants (33), suggesting that few genetic changes have occurred since *w*AlbB-I was first transferred to *Ae. aegypti* over 15 years ago (24). In 2015, the *w*AlbB-II variant from the *Ae. albopictus* strain UJU (origin Sulawesi, Indonesia) was transferred into an *Ae. aegypti* line with a Malaysian mitochondrial haplotype through microinjection (6). The donor *Ae. albopictus* UJU embryos carried a triple *Wolbachia* infection, comprising *w*AlbA, *w*Mel and *w*AlbB-II, but incomplete maternal transmission of the triple infection in *Ae. aegypti* allowed for isolation of a single-infection *w*AlbB-II line.

wAlbB-I and wAlbB-II populations were backcrossed regularly to natively uninfected populations from Australia and Malaysia respectively to control for genetic background. Uninfected populations were created through antibiotic treatment and the different combinations of nuclear background (Australian or Malaysian), mitochondrial haplotype (Australian or Malaysian) and *Wolbachia* infection status (wAlbB-I, wAlbB-II or uninfected) were generated through reciprocal backcrosses as explained in Ross et al. (2021). Two replicate populations of each combination were created and maintained separately. Both of these were included in *Wolbachia* density and quiescent egg viability measurements, while a single replicate population was tested for *Wolbachia* density under heat stress.

### *Wolbachia* detection and density

qPCR assays were used to confirm the presence or absence of *Wolbachia* infection and measure relative density. Genomic DNA was extracted using 250 μL of 5% Chelex 100 Resin (Bio-Rad laboratories, Hercules CA) and 3 μL of Proteinase K (20 mg/mL) (Roche Diagnostics Australia Pty. Ltd., Castle Hill New South Wales, Australia). Tubes were incubated for 30 minutes at 65°C then 10 minutes at 90°C. *Wolbachia* density was quantified with qPCR via the Roche LightCycler 480. Two primer sets were used to amplify markers specific to mosquitoes (forward primer mRpS6_F [5’- AGTTGAACGTATCGTTTCCCGCTAC-3’] and reverse primer mRpS6_R [5’-GAAGTGACGCAGCTTGTGGTCGTCC-3’]), and *w*AlbB (*w*AlbB_F [5’-CCTTACCTCCTGCACAACAA-3’] and *w*AlbB_R [5’-GGATTGTCCAGTGGCCTTA-3’]). For mosquitoes carrying the *w*Mel infection, *Wolbachia* density was determined using w1 primers (w1_F [5’-AAAATCTTTGTGAAGAGGTGATCTGC-3’] and w1_R [5’-GCACTGGGATGACAGGAAAAGG-3’], Lee et al. (2012)). Relative *Wolbachia* densities were determined by subtracting the Cp value of the *Wolbachia*-specific marker from the Cp value of the mosquito-specific marker. Differences in Cp were averaged across 2-3 consistent replicate runs, then transformed by 2^n^.

### Quiescent egg viability

We measured quiescent egg viability in *Ae. aegypti* populations with different combinations of *w*AlbB infection type (*w*AlbB-I, *w*AlbB-II or uninfected), mitochondrial haplotype (Au or My) and background (Au or My). Six cups filled with larval rearing water and lined with sandpaper strips were placed inside cages of blood fed females from each population. Eggs were collected five days after blood feeding, partially dried, then placed in a sealed chamber with an open container of saturated potassium chloride (KCl) solution to maintain a constant humidity of ∼84%. When eggs were 1, 4, 7, 10, 13 and 16 weeks old, small sections of each sandpaper strip were removed and submerged in water with a few grains of yeast to hatch. Four to six replicate batches of eggs were hatched per replicate population at each time point, with 40-125 eggs per batch. Hatch proportions were determined by dividing the number of hatched eggs (with a clearly detached egg cap) by the total number of eggs per female.

### *Wolbachia* density following heat stress

We measured *Wolbachia* density in adults after being exposed to cyclical heat stress during the egg stage. Eggs were collected from *Wolbachia*-infected populations (one replicate population each from *w*AlbB-I Au/Au, *w*AlbB-I Au/My, *w*AlbB-II My/Au, *w*AlbB-II My/My and *w*Mel). Four days after collection, batches of 40-60 eggs were tipped into 0.2 mL PCR tubes (12 replicate tubes per population) and exposed to cyclical temperatures of 29-39°C for 7 d in Biometra TProfessional TRIO 48 thermocyclers (Biometra, Göttingen, Germany) according to Ross et al. (2019). Eggs of the same age from each population were kept at 26°C. Eggs held at 29-39*°*C and 26°C were hatched synchronously and larvae were reared at a controlled density (100 larvae per tray of 500 mL water). Pupae were sexed and 15 males and 15 females per population and temperature treatment were stored in absolute ethanol within 24 hr of emergence for *Wolbachia* density measurements (see *Wolbachia* detection and density).

### Statistical analysis of density and phenotypic traits

Experimental data were analyzed using SPSS Statistics version 24.0 for Windows (SPSS Inc, Chicago, IL). Quiescent egg viability and *Wolbachia* density data were analyzed with general linear (mixed effect) models (GLMs). Replicate populations were pooled for analysis when effects of replicate population exceeded a P-value of 0.1 in prior analyses. Data for each sex were analyzed separately. For *Wolbachia* density, untransformed data (i.e. differences in Cp between *Wolbachia* and mosquito markers, before 2^n^ transformation) were used for analyses. We ran additional GLMs on *Wolbachia* density in each nuclear background separately due to significant interactions between background and *w*AlbB variant. For comparisons of *Wolbachia* density at different temperatures, we included temperature treatment (26 or 26-39°C) as a factor. We were unable to perform direct comparisons between *w*Mel and *w*AlbB strains due to using different markers for each strain; we therefore excluded *w*Mel from the overall analysis. We ran separate GLMs for each *w*AlbB variant due to significant two-way interactions. For quiescent egg viability, hatch proportions differed substantially between *w*AlbB-infected and uninfected populations. We therefore ran separate GLMs for *w*AlbB-infected and uninfected populations, with egg storage duration included as an additional factor for this trait. Replicate population (nested within *Wolbachia* infection status) was included as a random factor due to significant effects of replicate population for this trait. Squared egg storage duration was also included as a factor in the GLM due to the non-linear relationship between egg hatch proportion and storage duration in these populations.

### Multiplex PCR reaction

DNA was extracted from a pool of five female mosquitoes by crushing tissues in 200 μL of STE buffer. Each sample was then treated with 2 μL of Proteinase K (20 mg/mL) at 65°C for 30 min followed by a 10 min incubation step at 95°C. Tissue debris were removed by centrifugation for 2 min at 1,000 g and the supernatant was diluted 1/5 in water before PCR. Primer pairs specific of each *w*AlbB variant were designed to amplify a target gene present in one variant and absent in the other one. Primers were designed on an AAA-family ATPase protein (DEJ70_04410; forward: 5’-ATGTCTGTTTCTGCGTCTTG-3’; reverse: 5’-ATCGTCTTTATCCAGCCCAG-3’; 303 bp product) for *w*AlbB-I and on the phage tail formation protein I for *w*AlbB-II (WP_015587732.1; forward: 5’- AGAAATACTGCGCTGGGTAA-3’; reverse: 5’-GGATTGCTACATCTAGGCGA-3’; 497 bp product). As a DNA extraction control, primers were also designed to amplify the mosquito 18S gene (forward: 5’-CCCAGCTGCTATTACCTTGA −3’; reverse: 5’- TAAGCAGAAGTCAACCACGA-3’; 752 bp product). The three primer pairs were pooled in a multiplex PCR reaction using the Q5 High-Fidelity DNA Polymerase (New England Biolabs) in a 25 μL final volume as follows: 5 μL of buffer, 0.5 of 10 mM dNTPs, 1.25 μL of each 10 μM primer, 0.25 μL of DNA polymerase, 9.75 μL of water and 2 μL of DNA template. The PCR cycle used was: 98°C for 30s, 35 cycles of 10s denaturation at 98°C – 30s of annealing at 64°C – 1 min extension at 72°C, 2 min final extension at 72°C. PCR product were ran on a 1% agarose gel electrophoresis.

### Data availability

The *w*AlbB-II draft genome and raw sequencing data have been deposited at the NCBI GenBank database under the BioProject accession PRJNA800254 (assembly: JAKLOR000000000; Illumina reads: SRR17831854-SRR17832810; Oxford Nanopore reads: SRR17832811).

## Acknowledgments

The study was supported by Wellcome Trust (202888, 108508) to SPS and by the National Health and Medical Research Council (1132412, 1118640 [https://www.nhmrc.gov.au]) to AAH. LT and ASF were funded by the MRC (MC_UU_12018/12). The funders had no role in study design, data collection and analysis, decision to publish, or preparation of the manuscript.

## Supplemental Material

**Figure S1. Sequencing depth plots.** Mean sequencing depth per 500 bp window of Illumina reads mapped onto (A) *w*AlbB-II, (B) *w*AlbB-I, (C) *w*AlbB-FL2016 and (D) *w*AlbB-HN2016 gemomes.

**Figure S2. Maximum likelihood phylogenies of horizontally-transferred translocase subunit SecA genes.** (A) WP_019236968.1 and (B) WP_019236969.1 amino acids were aligned with their homologues and conserved sites (Gblocks) were used to build the phylogenies with PhyML. Node labels are bootstrap supports calculated from 100 replications.

**Table S1. List of genes being absent in at least one wAlbB genome.**

**Table S2. List of SNPs between wAlbB genomes.**

**Table S3. Statistical analysis of differences in density between *w*AlbB variants.**

**Table S4. Statistical analysis of *w*AlbB variant densities under heat stress.**

**Table S5. Statistical of quiescent egg viability.**

